# Sequencing of the human IG light chain loci from a hydatidiform mole BAC library reveals locus-specific signatures of genetic diversity

**DOI:** 10.1101/006866

**Authors:** Corey T. Watson, Karyn Meltz Steinberg, Tina A. Graves, Rene L. Warren, Maika Malig, Jacqueline Schein, Richard K. Wilson, Robert A. Holt, Evan E. Eichler, Felix Breden

## Abstract

Germline variation at immunoglobulin gene (IG) loci is critical for pathogen-mediated immunity, but establishing complete reference sequences in these regions is problematic because of segmental duplications and somatically rearranged source DNA. We sequenced BAC clones from the essentially haploid hydatidiform mole, CHM1, across the light chain IG loci, kappa (IGK) and lambda (IGL), creating single haplotype representations of these regions. The IGL haplotype is 1.25Mb of contiguous sequence with four novel V gene and one novel C gene alleles and an 11.9kbp insertion. The IGK haplotype consists of two 644kbp proximal and 466kbp distal contigs separated by a gap also present in the reference genome sequence. Our effort added an additional 49kbp of unique sequence extending into this gap. The IGK haplotype contains six novel V gene and one novel J gene alleles and a 16.7kbp region with increased sequence identity between the two IGK contigs, exhibiting signatures of interlocus gene conversion. Our data facilitated the first comparison of nucleotide diversity between the light and IG heavy (IGH) chain haplotypes within a single genome, revealing a three to six fold enrichment in the IGH locus, supporting the theory that the heavy chain may be more important in determining antigenic specificity.

## Introduction

Antibodies are essential components of the immune system that play key roles in processes associated with innate and adaptive immunity ^1^. They are expressed by B-cells as either cell surface receptors or secreted proteins, and are formed by two pairs of identical “heavy” and “light” immunoglobulin (IG) protein chains, encoded by genes located at three primary loci in the human genome: the IG heavy chain (IGH) at 14q32.33, and the two IG light chain regions, lambda (IGL) and kappa (IGK), located at 22q11.2 and 2p11.2 ^2^. Specifically, through a unique mechanism referred to as V-(D)-J recombination ^3^, individual Variable (V), Diversity (D), and Joining (J) genes at the IGH locus, and V and J genes at either the IGK or IGL loci recombine somatically at the DNA level to generate templates for the subsequent transcription and translation of antibody heavy and light chains, respectively. V-(D)-J recombination is accompanied by the random addition and deletion of nucleotides at the junctions of the combined V, D, and J genes by terminal deoxynucleotide transferase (TdT). The extreme variability observed in expressed antigen-naïve B-cell antibody repertoires is due to this combinatorial and junctional diversity, and partly ensures that the immune system is able to recognize and mount effective immune responses against a diverse range of potential pathogens. At the population and species level, IG haplotype and allelic variation also make important contributions to the diversity of expressed antibody repertoires ^4–7^; however, broader roles of IG genetic polymorphism in antibody function are generally less well understood ^8^.

Over the past three decades, extensive catalogues of genetic polymorphisms have been established for human IGH, IGK, and IGL genes (IMGT; the international ImMunoGeneTics information system; www.imgt.org ^2,9^). Importantly, not only has IG genetic diversity recently been implicated in inter-individual variation in expressed antibody repertoires ^4–6^, but genetic variants in both IG coding regions and well-characterized regulatory motifs are also known to influence antibody expression and function, and mediate risk of disease phenotypes ^10–12^. However, our understanding of germline variability at the IGH, IGK, and IGL loci remains severely limited, especially in terms of haplotype structure (*i.e*., large segmental duplications and deletions) as well as coding and non-coding sequence polymorphisms. We have begun to uncover this variability in the IGH locus, leading to complete nucleotide resolution descriptions of large structural variants (insertions, deletions, duplications, and complex rearrangements), including novel functional IGHV genes and alleles ^7^.

In the present study we use the same approach as our previous characterization of IGH to analyze complete haploid reconstructions of the IGK and IGL loci, which to date, compared to IGH, remain less well investigated. The IGL region has only been fully sequenced and analyzed once in its entirety ^13^, using multiple cosmid and BAC library resources. The region spans approximately 0.9 MB, and includes 69 IGLV, seven IGLJ, and seven IGLC functional/open reading frame (ORF) genes and pseudogenes. Additional V, J, and C genes not present in this haplotype are known to occur as insertion variants in the human population ^2,14–16^. Similarly, the initial sequence of the IGK locus was generated from a composite assembly of cosmid, bacteriophage, and BAC clone libraries ^17^. A unique feature of the IGK locus is that it includes two large inverted segmental duplications (SDs) that comprise distinct V-gene containing regions (termed proximal and distal); these regions remain separated by a large, currently unsequenced, assembly gap. The proximal region spans 0.54 MB and includes 69 IGKV functional/ORF genes and pseudogenes, whereas the distal region spans 0.43 MB and includes 62 V genes and pseudogenes (distal V genes are denoted by a “D”; *e.g.*, *IGKV1D-13*), five functional IGKJ genes, and a single functional IGKC gene also reside downstream of the proximal V gene cluster.

Haplotypes spanning the IGK proximal region, and a portion of the distal region have also been sequenced using BAC clones from the RPCI-11 library ^18^. Both IGL and IGK are known to exhibit V, J, and C gene allelic and structural variation ^2,14–16,19–21^.

As a means to improve existing genomic resources in the IG loci, we have sequenced the IGK and IGL gene clusters from the CHORI-17 BAC library (CH17, BACPAC resources), previously constructed from a haploid hydatidiform mole cell line (CHM1htert). Together with data produced previously for the IGH locus from the CH17 library ^7^, these sequences represent the first complete human haplotypes of all three IG loci from a single individual. Using these data, we have conducted the first comparison of genetic diversity between the three IG regions in the same haploid genome. Our analyses have facilitated the identification of novel single nucleotide polymorphisms (SNPs) within IGL and IGK gene coding regions and regulatory elements, and also revealed a novel large sequence conversion event between the IGKV proximal and distal regions. In addition, our unbiased tri-locus comparison shows a striking enrichment of structural and nucleotide diversity in the IGH locus, confirming previous suggestions that germline variation within the light chain gene loci is lower than that observed in IGH ^15,22,23^.

## Results

### IGL and IGK reference sequences from the CH17 BAC library

We analyzed sequences from 17 CH17 BAC clones (IGL, 9; IGK, 8) comprising tiling paths across the two loci (Figures 1 and 2) -- one of the clones in IGK, CH17-198L18, had been sequenced previously. Clones unique to either IGK or IGL were then used to construct locus-wide contigs; clones in IGKV proximal and distal regions were aligned separately because the gap separating these two regions was not completely filled by the current sequencing effort.

**Figure 1.**
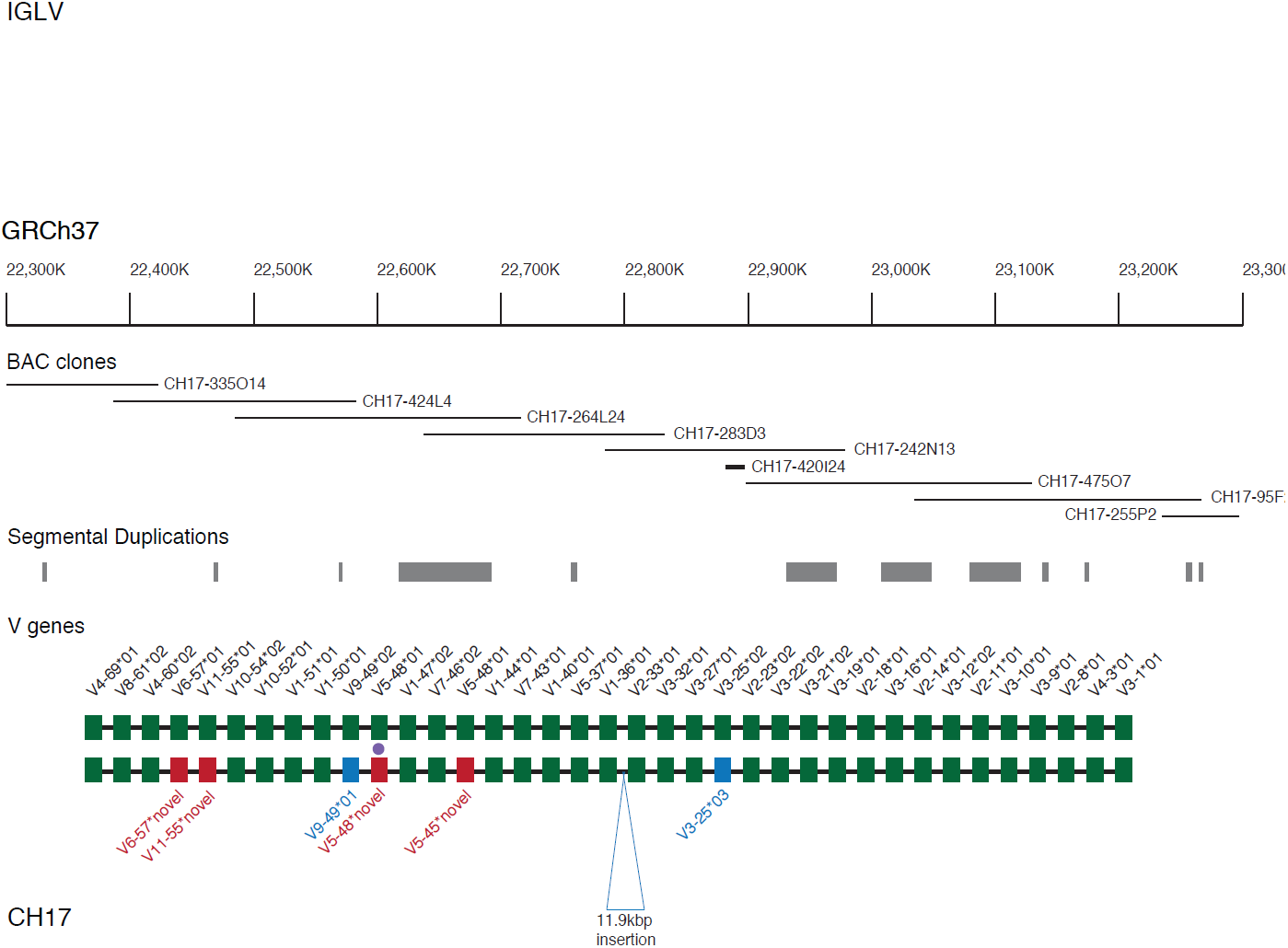
IGLV gene comparison between CH17 and Kawasaki haplotypes. Tiling path of sequenced CH17 BAC clones and functional and ORF IGLV genes annotated on GRCh37 and CH17 are depicted by filled boxes, with corresponding locus and allele identifiers located above and below the haplotypes. Genes/alleles shared between haplotypes are indicated by filled green boxes. Genes and alleles that are unique to CH17 (not present in the NCBI reference/Kawasaki haplotypes) are indicated by boxes with other colors (red for non-synonymous and blue for synonymous). Filled purple circles denote novel alleles with polymorphisms resulting in a nonsense mutation (pseudogene). The 11kbp insertion in the CH17 IGL path is indicated.

**Figure 2.**
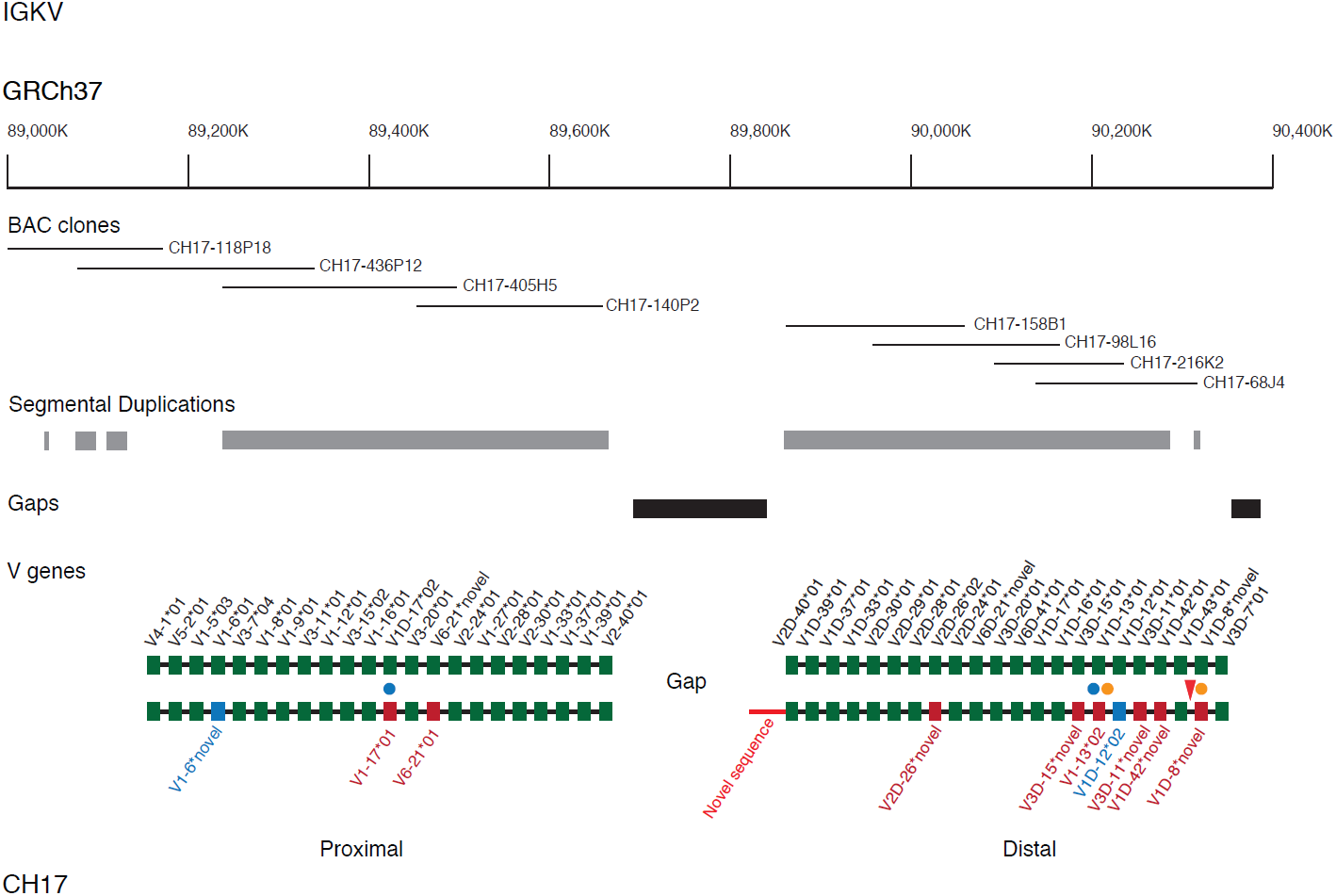
IGKV gene comparison between CH17 and Kawasaki haplotypes. Tiling path of sequenced CH17 BAC clones and functional and ORF IGKV genes annotated on GRCh37 and CH17 are depicted by filled boxes, with corresponding locus and allele identifiers located above and below the haplotypes. Genes/alleles shared between haplotypes are indicated by filled green boxes. Genes and alleles that are unique to CH17 (not present in the NCBI reference/Kawasaki haplotypes) are indicated by boxes with other colors (red for non-synonymous and blue for synonymous). Filled blue circles denote loci at which proximal or distal alleles were observed at the alternate locus (*e.g.*, proximal allele at distal locus); and orange circles denote allelic differences between the haplotypes with respect to regulatory elements. The novel sequence extending into the gap in the CH17 IGK path is shown in red. The 21bp indel polymorphism upstream of IGKV1D-8 is indicated with a red arrow.

For IGL, a single 1.25 Mbp contig was generated from 9 BACs spanning 175.7 Kbp upstream of the functional IGLV gene, *IGLV4-69*, to 191.3 Kbp downstream of *IGLC7* (Supplementary Figure 1). In total, we identified 37 of the 38 known functional/ORF IGLV genes, seven functional J genes, and four functional C genes. The remaining IGLV gene, IGHV5-39 was not found in CH17, consistent with it being an insertion polymorphism ^15,21^. Sequence comparisons within IGLV, IGLJ, and IGLC genes revealed allelic differences between CH17 and the Kawasaki haplotype at six V genes, two J genes, and one C gene (Figure 1; Supplementary Table 1). Six of these alleles, five of which were novel (*IGLV6-57*, *IGLV11-55*, *IGLV5-45*, *IGLV5-48*, and *IGLC7*), included non-synonymous changes. Notably, the novel allele identified at *IGHV5-48* included a nonsense mutation that introduced a premature stop codon in the framework 3 region of the protein; 15 additional SNPs were also characterized within the exons of this gene. Prior to this study, only a single allele of *IGLV5-48* had been described, classified as an ORF due to an uncharacteristic single nucleotide difference in the heptamer portion of the recombination signal (RS: <u>T</u>ACAGT instead of <u>C</u>ACAGTG ^24^). SNPs characterized in the remaining four novel functional/ORF alleles were represented in the 1000 genomes project (1KG) dataset ^25^. Previously described regulatory motifs ^13^, including RS sequences, associated with each of the 37 identified functional/ORF IGLV genes were also inspected for previously uncharacterized variants in the CH17 haplotype, but no SNPs in these regions were identified.

Eight BAC clones were analyzed in the IGK locus, forming two independent contigs, one in the proximal region totalling 644 Kbp, and a second in the distal region of the locus totalling 466 Kbp of contiguous sequence (Figure 2). The proximal contig, containing four BACs, spanned from 170 Kbp downstream of *IGKC* to within the intron of *IGKV2-40*; thus, this contig lacked 10.9 Kbp of known sequence upstream of *IGKV2-40* characterized in the initial genomic description of the locus, representing a small gap in the CH17 sequence. In the distal region, four complete BACs were assembled into a single contig spanning 60 Kbp upstream of *IGLV2D-40* to 22 Kbp downstream of the most distal gene in the locus, *IGHV3D-7*. This sequence included ∼49 kbp of additional sequence (compared to the Kawasaki haplotype) extending into the assembly gap between the proximal and distal units, which is predicted to be 800 kbp ^17,26^. The sequence extending into the unsequenced gap is dominated by complex repeats (Supplementary Figure 1), likely contributing to the difficulty of completing assemblies in this region. Alleles at forty-four functional/ORF IGKV genes, 22 in each of the proximal and distal regions, as well as five IGKJ genes, and a single IGKC gene in the proximal region were characterized and compared to those found in the IGK Kawasaki haplotype (Figure 2; Supplementary Table 2). We observed ten allelic differences at IGKV gene loci, and one allelic variant at *IGKJ2*; ten of these allelic variants, including *IGKJ2*04*, involved non-synonymous changes. We characterized nine novel alleles that were not represented in IMGT, including three that were observed in the Kawasaki haplotype. In two instances, we observed the presence of alleles that had been previously classified as either “distal” or “proximal” alleles residing at loci in the alternate location. For example, we found an allele that matched with 100% sequence identity to *IGKV1-13*02* at the *IGKV1D-13* locus in the CH17 haplotype, for which there had previously been only one allele described, *IGKV1D-13*01*. *IGKV1D-13*01*, which differs from *IGKV1-13*02* by only a single nucleotide, was initially classified as an ORF due to an abnormal V-heptamer sequence, but was later identified in a productive rearrangement, indicating that it is likely functional ^27^.

**Table 1.** IGK and IGL loci CH17 BAC clones and contig statistics

| **Locus** | **Size Bp** | **Bp overlap with subsequent clone** | **Bp Mismatch** | **Genbank Accession** |
| --- | --- | --- | --- | --- |
| **IGK PROXIMAL** |  |  |  |  |
| CH17-118P18 | 204,603 | 129,222 | 0 | AC244205 |
| CH17-436P12 | 203,033 | 25,144 | 1 | AC243970 |
| CH17-405H5 | 255,429 | 63,619 | 1 | AC245015 |
| CH17-140P2 | 198,999 | NA | NA | AC244255 |
| **IGK DISTAL** |  |  |  |  |
| CH17-158B1 | 211,421 | 134,551 | 1* | AC233264** |
| CH17-98L16 | 214,723 | 59,786 | 0 | AC245506 |
| CH17-216K2 | 202,759 | 142,980 | 1* | AC243981 |
| CH17-68J4 | 181,341 | NA | NA | AC247037 |
| **IGL** |  |  |  |  |
| CH17-335O14 | 213,458 | 69,894 | 0 | AC245452 |
| CH17-424L4 | 192,598 | 63,215 | 0 | AC245517 |
| CH17-264L24 | 235,497 | 62,738 | 0 | AC245060 |
| CH17-238D3 | 189,776 | 79,534 | 1 | AC245291 |
| CH17-242N13 | 213,099 | 74,994 | 1 | AC246793 |
| CH17-420I24 | 210,075 | 170,533 | 0* | AC244250 |
| CH17-475O7 | 206,505 | 50,665 | 0 | AC244157 |
| CH17-95F2 | 196,325 | 41,691 | 1 | AC245028 |
| CH17-355P2 | 215,750 | NA | NA | AC245054 |

\*Bp mismatches do not include differences involving microsatellite repeat sequence; \*\* Previously sequenced; NA, not applicable

**Table 2.** IG region genomic feature statistics and SNP density comparisons

| **GENOMIC FEATURE** | **IGHV** | **IGLV** | **IGKV** |
| --- | --- | --- | --- |
| Total bp Comp | 834,802 | 760,133 | 858,805 |
| Total SNPs | 2,897 | 854 | 519 |
| Seg Dup fraction ($\geq 90\%$) | 0.377 | 0.278 | 0.304(0.887)<sup>1</sup> |
| Seg Dup fraction (90-95%) | 0.138 | 0.2 | 0.287(0.287)<sup>1</sup> |
| Seg Dup fraction (95-100%) | 0.288 | 0.146 | 0.051(0.826)<sup>1</sup> |
| Repeat coverage | 0.534 | 0.433 | 0.469 |
| # of functional/ORF V gene loci<sup>2</sup> | 42 | 37 | 44 |
| **SNP DENSITY** | **IGHV** | **IGLV** | **IGKV** |
| locus-wide | 0.0035 | 0.0011 | 0.0006 |
| F/ORF V gene | 0.0039 | 0.0015 | 0.0017 |
| Non Seg Dups | 0.0031 | 0.0012 | 0.0004 (0.0004)<sup>1</sup> |
| Seg Dups | 0.0042 | 0.001 | 0.001 (0.0006)<sup>1</sup> |
| Seg Dups (90-95%) | 0.0033 | 0.001 | 0.001 (0.001)<sup>1</sup> |
| Seg Dups (95-100%) | 0.0045 | 0.0014 | 0.001 (0.0006)<sup>1</sup> |
| Repeats | 0.0031 | 0.0011 | 0.0006 |

<sup>1</sup>Calculations in parentheses include inverted segmental duplications between the proximal and distal units. <sup>2</sup>Only those genes found in the CH17 and reference haplotypes were considered, excluding duplications and deletions described in IGHV (Watson *et al.*, 2013).

Interestingly, from our analysis of regulatory elements, we found that the allele in the CH17 haplotype at this locus (*IGKV1-13*02*) was associated with a typical, non-mutated V-heptamer sequence. Thus, this functional V-heptamer variant may also be associated with *IGKV1D-13* alleles (*e.g.*, *IGKV1D-13*01* noted above), and facilitate their expression.

Discrepancies have also been noted regarding the functionality of *IGKV1-8*, for which only one coding allele is known. Genomic descriptions of this gene have revealed a 21 bp deletion in an upstream regulatory element, which had been predicted to disruption of promoter function and inhibit expression (mutation observed in the Kawasaki haplotype^28^); however, *IGKV1-8*01* has been shown to be expressed in some cases ^27^. Potentially explaining this discrepancy, we found that the previously described *IGKV1-8*01* 21 bp promoter deletion was not present in the CH17 haplotype, suggesting that this germline indel variant could contribute to variation in the expression of alleles at this locus.

### Characterization of structural variants in IGL and IGK CH17 haplotypes

A direct comparison of the IGL CH17 and Kawasaki haplotypes revealed the presence of only a single structural variant. This 11.9 Kbp insertion was located in the region between the psuedogenes *IGLV7-35* and *IGLV2-34* within the BAC, CH17-242N13; the region between these pseudogenes spans ∼120 Kbp and is devoid of IG genes. Gene prediction analysis did not identify any genes within the insertion, nor did the insertion disrupt the non-IG related genes, *ZNF280B*, *ZNF280A*, and *PRAME*, located in this region; the breakpoints of the event occurred between *ZNF280A* and *PRAME*. No structural variants were observed in the IGK CH17 haplotype.

We also searched the CH17 haplotypes for all IGL and IGK gene/allele sequences classified as “not located” in the IMGT database, meaning that these genes have not been located within IGL or IGK loci. Using this approach we mapped the pseudogene *IGLV2-NL1* to the locus in CH17 corresponding to the position of the pseudogene *IGLV2-34* in the Kawasaki haplotype; *IGLV2-NL1* matched CH17 at this locus with 100% sequence identity, suggesting that *IGLV2-NL1* and *IGLV2-34* genes are allelic rather than distinct loci. No additional exact matches between the CH17 haplotype and other “not-located” V genes were observed.

### Analysis of proximal and distal regions in CH17 and Kawasaki haplotypes reveals evidence for sequence exchange

Previous analysis of sequence similarity between shared homology blocks of proximal and distal segmental duplication units, which comprise the majority of the IGK locus, revealed that these two regions are >98% similar over most of the region ^17^. Segmental duplications are known to facilitate sequence exchange via non-allelic homologous recombination and interlocus gene conversion ^29,30^; however, given the lack of reference sequence data this has not been investigated in the IGK locus. To address this, we conducted pair-wise comparisons of distal and proximal regions from CH17 and Kawasaki haplotypes to search for large tracts of shared sequence (Figure 3A). The expectation is that sequence should be most similar between homologous regions in the alternate haplotype, whereas higher similarity between proximal and distal regions within a haplotype would suggest the occurrence of sequence exchange. Using this approach, we identified a large ∼16.7 Kbp region that showed higher identity between the proximal and distal units of the Kawasaki haplotype than between the CH17 distal and Kawasaki proximal units (Figure 3A). This region included two IGKV genes for which we observed allelic variants between the Kawasaki and CH17 haplotypes. Four-way sequence alignments of this region show that the CH17 distal unit was most unique compared to the other three sequences (Figure 3B), providing evidence that distal and proximal units have undergone sequence exchange, making the distal unit more similar to the proximal unit in this region in the Kawasaki haplotype. It is important to note, however, that the Kawasaki distal fragment harbors many unique bp differences compared to the other three sequences (blue tick marks, Figure 3B), which could be suggestive of the occurrence of mutation following the predicted sequence exchange event. Further analysis of this multi-sequence alignment using the DSS method for recombination detection also predicted two potential flanking recombination breakpoints within the expected regions based on visual inspection of the sequence alignment and comparison of sequence similarities. We also analyzed sequence from two BAC clones in the proximal and distal clusters from the RPCI-11 BAC library ^18^; this analysis revealed that these carried the same variants observed in the Kawasaki haplotype.

**Figure 3.**
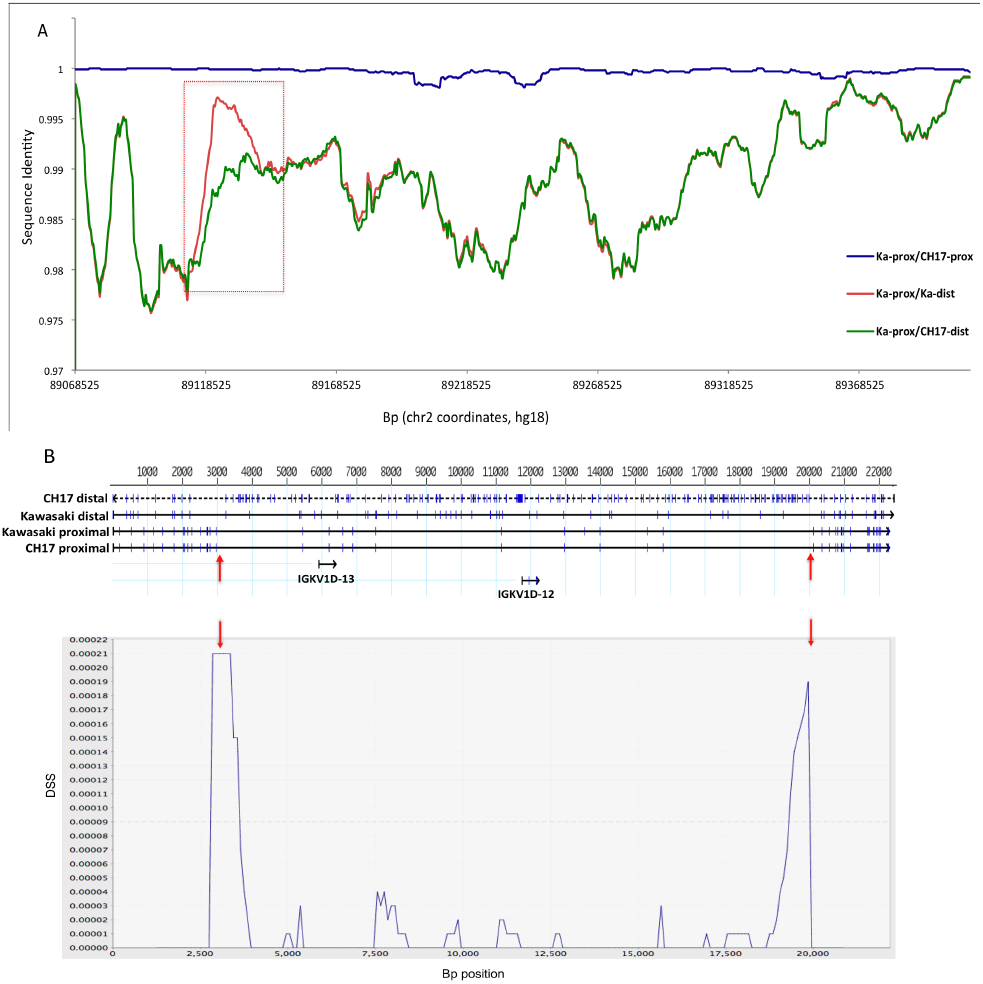
Detection of putative sequence exchange event between IGK proximal and distal regions. (A) Pair-wise alignments between proximal and distal segmental duplications in the CH17 and Kawasaki haplotypes (Abbreviations: Ka, Kawasaki; prox, proximal; dist, distal). The region where Ka-prox and Ka-dist show stronger similarity than CH17-dist and Ka-prox highlights a potential region of sequence exchange between the two CH17 units (red box). (B) Top panel shows a four-way sequence alignment of a 22.5 Kbp region from the proximal and distal units from within the red box in (A). Blue tick marks indicate bp SNP differences between the sequences. Upward pointing red arrows indicate boundaries of regions where the Kawasaki distal sequence aligns with a higher sequence similarity to the Kawasaki and CH17 proximal sequences than to the CH17 distal sequence, indicative of exchange between proximal and distal regions of the Kawasaki haplotype (sequence similarities: Ka-dist/CH17-dist=98.7%; Ka-dist/Ka-prox=99.7%). A DSS recombination analysis (McGuire and Wright, 2000; see methods) using the same four-way sequence alignment is shown in the bottom panel. The two peaks with the strongest DSS values (downward pointing red arrows) correspond to the predicted breakpoints shown in the top panel based on sequence similarity values. The dotted line across the chart indicates the significance threshold based on the null distribution of DSS values calculated assuming no recombination.

### SNP diversity and genomic features in IGHV, IGLV, and IGKV gene regions

Compared to the number of allelic variants observed between the IGHV CH17 haplotype and the reference genome (19 allelic variants/40 V gene loci^31^), V gene allelic variation described in this study for IGL (6 allelic variants/37 V gene loci) and IGK (10 allelic variants/44 V gene loci) was noticeably lower. This prompted us to also compare other genomic characteristics between the three loci. Excluding regions of structural variation between haplotypes, we first generated SNP calls (not including gaps and single bp indels) between the CH17 and reference haplotypes for all three loci; 491, 1046, and 2897 SNPs were identified for IGKV, IGLV, and IGHV, respectively (Table 2; Figure 4, left panel). After cross referencing these SNPs with dbSNP135 and 1KG datasets, 74, 110, and 407 SNPs in the IGKV, IGLV, and IGHV gene regions were determined to be novel variants, not represented in either dataset. Not surprisingly, given the number of SNPs in the 1KG datasets, fewer SNPs at each locus were represented in dbSNP (Figure 4). We examined these sites in publicly available Illumina data generated from the CHM1 genomic DNA to determine if the novel SNPs were supported by an orthogonal platform. We identified 73/74, 103/110, and 406/407 sites in IGKV, IGLV, and IGHV, respectively, that are supported by the Illumina data. The discrepancies may represent sequencing errors. The novel SNPs for each region reported in GRCh37 coordinates are in Supplementary Table 3.

**Figure 4.**
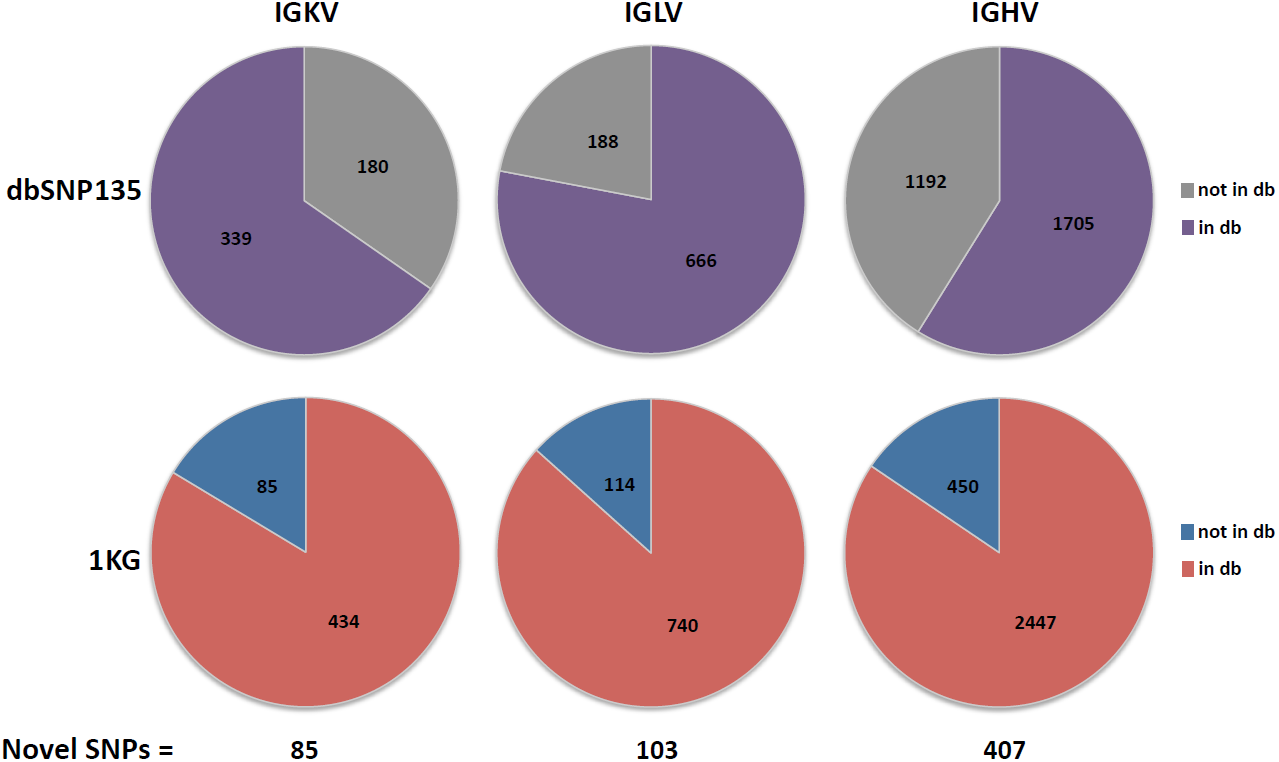
Representation of SNPs identified in the CH17 IG V gene regions in public SNP databases. The number of SNPs identified in IGKV, IGLV, and IGHV gene regions based on alignments of CH17 to Matsuda and Kawasaki haplotypes (excluding gaps). The fraction of SNPs represented in dbSNP135 (top) and 1KG (bottom) databases (db) are shown, the total number of novel SNPs not found in either database is indicated at the bottom of the left panel.

Consistent with observations based on V gene allelic variation, SNP density in IGHV (0.0035) was approximately 3-fold higher than in IGLV (0.0012) and 6-fold higher than in IGKV (0.0006); SNP densities were slightly elevated within functional/ORF V genes in each region compared to the calculated locus-wide values (Table 2). An analysis of genomic features in the three loci also showed that the fraction of each locus covered by repeat content was highest in IGHV, whereas that covered by SDs, not surprisingly was highest in IGKV (Table 2). However, when SDs associated with the IGKV proximal/distal duplication event were excluded, SD coverage in IGHV (37.7%) was found to be much higher than both of the light chain loci (IGLV, 24.6%; IGKV, 28.2%; Table 2). When only SDs exhibiting > 95% sequence identity were considered, this difference was even more striking (IGHV, 28.8%; IGLV, 12.9%; IGKV, 1.0%). Interestingly, in IGHV, SNP density was higher in regions of segmental duplication compared to regions not covered by SDs, especially true when considering only those regions covered by SDs with >95% sequence identity (Table 2); however, this difference was not found to be significant after permuting SD *vs.* non-SD region assignments and re-assessing SNP densities of 10,000 MCFDR samplings (*P* > 0.05). SNP density was also increased in segmentally duplicated regions of IGKV with >95% sequence identity compared to regions not covered by SDs, but again, this difference was not significant (*P* > 0.05).

Given the telomeric location of IGHV and the known differences in nucleotide substitution patterns within telomeres compared to the rest of the genome ^32^, we next assessed CHM1 SNP density within and around autosomal telomeres and centromeres. We found that mean SNP densities were ∼2-fold higher in telomeric regions (within 3 Mbp = 0.001; 1 Mbp = 0.0009) when compared to centromeric regions (within 3 Mbp = 0.0005; 1 Mbp = 0.0004; Supplementary Figure 2A). Interestingly, SNP density within 1 Mbp of the q arm telomere of chr14 harbouring IGHV (0.002) was higher than all other telomeric regions, >2-fold higher than the telomeric average (Supplementary Figure 2B). In contrast, when CHM1 SNP densities within 3 Mbp of all telomeres were compared, the region on the q arm of ch14 (including both IGHV and non-IGHV sequence) no longer stands out, suggesting that IGHV may have some unique properties contributing to higher than average genetic diversity. To place this in the context of the analysis conducted within the IG regions above, we also found that the q arm of ch14 has the second highest overlap with SDs (34%; Supplementary Figure 3).

### Discussion

We present data for the first haplotypes of the human IGL and IGK loci from the same haploid genome, representing only the second full-length references constructed for these regions to date. From the CH17 clones, 12 novel alleles were identified in the two loci, including four IGLV alleles, seven IGKV alleles, and one IGLC allele. Two recent assessments of IGK allelic variation -- one of a public dataset of 435 expressed sequences ^33^, and a second of deep-sequenced antibody repertoires from four individuals ^34^ -- concluded that, unlike IGH, IGK allelic datasets are likely to be mostly complete, as only two putative novel alleles were identified from these analyses ^33^. IGHV and IGKV repertoire sequencing in a single individual also supports these observations, finding that nearly 25% of characterized alleles in IGHV were novel, compared to 0% in IGKV ^35^. However, the fact that we identified 11 novel light chain V gene alleles from a single haploid genome implies that additional efforts to identify unreported alleles in IGL and IGK are warranted. Importantly, as noted previously in IGHV ^7^, SNPs associated with novel alleles identified in IGL and IGK were present in the 1KG dataset, further supporting the notion that the 1KG dataset could serve as a useful resource for future investigations of IG gene diversity and the identification of novel polymorphisms. However, given the prevalence of segmental duplication in the IG loci, it will undoubtedly be essential to consider the impact of paralogous sequence variants (PSVs) when assessing 1KG SNP data in these regions, as complex and duplicated sequence structure are known to confound SNP characterization ^36,37^.

In addition to novel allelic variants within IGL and IGK coding regions, variants involving regulatory elements of two IGKV genes (*IGKV1-8* and *IGKV1D-13*) were also identified. In both cases, regulatory elements previously associated with alleles at these loci were predicted to inhibit their expression; however, the alleles described for these genes in the CH17 haplotype were associated with regulatory region variants that would be expected to exhibit normal gene expression. The importance of such polymorphisms is that they can result in variable levels of gene expression, including the loss of genes/alleles from expressed repertoires. In addition to those noted above, several other IGKV genes are also classified as ORF genes based on irregular regulatory sequence motifs ^27^. Notably, *IGKV2D-29*02*, previously referred to as “VA2c”, has a non-canonical V-heptamer, but has also been shown to occur in productive rearrangements; thus, like *IGKV1-8* and *IGKV1D-13* alleles described in CH17, the expression of *IGKV2D-29*02* could be explained by a previously undescribed regulatory sequence variant. Interestingly, a third allele at the *IGKV2D-29* locus, “V2b”, has also been shown to have a defective RS V-heptamer, which results in decreased expression and has been implicated in susceptibility to *Haemophilus influenzae* type b infection ^10,38^.

Perhaps the most important contribution of the full IGK and IGL sequences of the CH17 haplotype data presented here is that, for the first time, locus-wide genetic diversity between IGH, IGL, and IGK could be compared in the same haploid genome in relation to corresponding reference haplotypes. Our first observation from this comparison was that the number of V gene allelic differences among the CH17 haplotypes was highest in IGHV. This finding is perhaps not surprising given that allelic richness is also known to be highest in IGHV based on available genetic data in the IMGT database. In addition, studies of substitution patterns in heavy and light chain V genes have also revealed evidence for increased diversity in IGHV compared to IGLV and IGKV ^23,39^. Strikingly, however, when we extended our comparison to include all SNPs within the three loci, we found that locus-wide genetic diversity was also much higher in IGHV, indicating that increased diversity in this locus is not limited to within V gene coding regions. IGHV SNP density is also higher than that observed for killer cell immunoglobulin-like receptor/leukocyte Ig-like receptor and T cell receptor alpha loci, calculated at ∼0.0016 for both regions (Steinberg *et al.*, *manuscript in preparation*).

Similar to patterns noted from V gene allelic diversity and substitution patterns, fewer SVs have also been reported in IGLV and IGKV compared to IGHV. In IGL, for example, only three insertion-deletion variants of functional genes have been identified, involving *IGLV1-50*, *IGLV8-61*, and *IGLV5-39* ^15,21^; although, the deletion of *IGLV8-61* was identified in only a single individual. Likewise, in the IGKV locus, aside from an identified rare haplotype containing a deletion of the entire distal IGKV gene cluster ^40,41^, only a single functional V gene insertion, including the gene *IGKV1-NL1*, has been identified ^19^. Limited evidence suggesting putative IGKV gene duplications/deletions (*e.g.*, *IGKV1-5* and *IGKV3-20*) has also more recently been noted from expressed antibody repertoire data, although these await confirmation as germline polymorphisms ^34^. Thus, in total, liberal estimates from the literature indicate that only 4-6 light chain functional/ORF V genes are known to occur in SVs, in stark contrast to ∼29 in IGHV ^7,8^. In line with this difference, four V gene-containing SVs were identified in the CH17 IGHV haplotype compared to zero defined in IGLV and IGKV.

Two factors suggest that the higher rate of SVs in IGH may be attributable to the increased fraction of the locus covered by segmentally duplicated sequences. First, segmental duplications are known to be associated with SVs genome-wide ^42^, and second, duplications have been shown specifically to facilitate structural variation in IGH ^7^. Segmental duplications and tandem repeats also mediate sequence exchange either through gene conversion or recombination, that can either result in the homogenization of paralogous sequences ^29,30,43^, or in an increase in genetic diversity ^44,45^. Illustrating the latter of these two scenarios, we found that for IGHV, SNP density was highest in regions including segmental duplications, particularly those with >95% sequence identity with their paralogs; similar trends were not noted for SNP density estimates calculated within non-SD repeat regions (Table 2). Importantly, SDs with >95% identity comprised nearly twice the fraction of IGHV sequence than IGLV sequence, which may in part explain the differences observed in SNP density between IGH and IGL. It is also worth noting that the difference in the fraction of SDs between IGH and the light chain regions would be greater if the 222 Kbp of novel sequence (comprised primarily of SDs) identified in IGH by our previous study ^7^ were also included. In addition to this, assessments of CHM1 SNP density across autosomal telomeres and centromeres revealed that the telomeric region on chr14 containing IGHV had both elevated levels of SNP diversity, and increased SD overlap, compared to analogous regions on other autosomes. This suggests that the genomic location of IGHV has also likely contributed to the increased genetic variation we observe in this locus compared to IGKV and IGLV.

The distinct clustering and genomic partitioning of V gene families within the IGL locus, and the observation that, compared to IGH and IGK, there are fewer IGL orphans present in other regions of the genome, has prompted the suggestion that IGL genes have undergone less “evolutionary shuffling” ^22^, which may be linked to lower levels of diversity in the locus ^15^ and would be consistent with the results presented here. In comparison to the other two loci, we found IGKV to have the lowest locus-wide SNP density, nearly 6-fold lower than that observed in IGHV. Due to the large inverted duplication of the proximal and distal regions, over 80% of the IGKV locus consists of SDs with >95% sequence identity. This suggests that, unlike in IGHV, SDs may be responsible for sequence homogenization rather than an increase in SNP diversity. The fact that we found evidence of a large tract of sequence exchange between the proximal and distal IGKV units lends support to this notion. However, fully understanding the relationship between segmental duplication and SNP density in the human IG regions will undoubtedly require further sequencing and comparisons of additional haplotypes. We must also acknowledge the potential for confounding effects related to the use of mosaic reference sequences for this comparison ^13,17,31^, which were generated from multiple large insert libraries constructed from diploid tissues, in some cases of unknown ethnicity. Considering this, it is possible that a comparison of CH17 IG regions to references generated from individuals with different ethnic backgrounds could result in artifactual differences between loci. However, because our findings are supported by existing V gene allelic variation data at the population level, it seems unlikely that the difference in variability between loci is due to ethnic origin of the tissue.

If the difference in SNP density observed here between the IG V gene clusters is in fact genuine, it raises the question of whether increased genetic diversity in IGH has any functional consequences. Given that SNP density within V gene coding regions in the CH17 haplotype was also higher in IGH compared to IGL and IGK, it could be speculated that mechanisms associated with an increased number of polymorphisms locus-wide in IGHV, could by default, result in greater IGHV gene diversity and a more variable expressed antibody repertoire. Intriguingly, in natural antibodies, the IG heavy chains are considered to play a more prominent role in epitope binding than IG light chains, although this is primarily attributed to residues of the third complementary determining region (CDRH3) not encoded by IGHV gene segments ^46^. However, there are several examples demonstrating essential functions of IGHV germline-encoded variation in antigen specificity; for example, residues encoded by germline *IGHV1-69* alleles have been shown to make important contributions to neutralizing antibody responses against influenza, hepatitis C, and Middle East Respiratory Syndrome coronavirus ^11,47,48^. Whether increased genetic diversity in IGHV is associated with the dominant role of the heavy chain in antibody function remains to be seen. In addition, there is still much to be learned about the contribution of IG genetic polymorphism to variability in expressed repertoires and the implications of this variation for susceptibility to infectious and autoimmune diseases, responses to therapeutic antibodies and vaccines, and other clinical outcomes. These outstanding questions stress the importance of accurately representing standing genetic variation in the human IG loci.

## Materials and methods

### Sequencing of IGL and IGK regions from the CHORI-17 BAC library

BAC-end reads from the CHORI-17 hydatidiform mole BAC library mapping to the GRCh37 reference genome were used to identify and select clones within the two light chain regions. The IGL and IGK genes are located at distinct loci in the genome, 22q11.2 and 2p11.2, respectively ^2,13,17,49–51^. Nine clones mapping to the IGL region, and eight clones mapping to IGK were picked for complete sequencing. As described in Watson et al^7^ clones were shotgun sequenced using high quality capillary-based Sanger sequencing and assemblies were constructed and finished on a per clone basis. Fully assembled overlapping BAC clones were then used to create contiguous assemblies spanning the two IG light chain regions using SeqMan Pro (DNA Star, Lasergene, Wisconsin, USA).

### Annotation of V, J, and C genes and regulatory regions from BAC clones

Sequences of all functional and open reading frame (ORF) C, J, and V genes (based on IMGT classification) were downloaded from IMGT and *Vega* databases (www.imgt.org, vega.sanger.ac.uk). All sequences were aligned to the completed contigs of each locus using SeqMan Pro, the positions of which were confirmed using BLAST ^52^. Sequences corresponding to each of the mapped C, J, and V genes were extracted from the CH17 contigs, and alleles at each locus were assigned using IMGT V-QUEST ^53,54^. “Novel” alleles were defined as those not found in the IMGT database. To search for potential variants in previously characterized regulatory sequences, SNPs determined from alignments of the CH17 haplotype and references generated previously by Kawasaki *et al*. (we refer to these as “Kawasaki haplotypes” ^13,17^) were tested for overlap with 250 bp regions immediately up and down stream of functional and ORF gene exons. Exon coordinates as determined by *Vega* annotations for each gene were downloaded from UCSC (www.genome.ucsc.edu), and SNP/gene region overlap was assessed using BEDTools version 2.1 ^55^. For those genes in which a SNP was found to occur within the defined regions, sequences in question from the CH17 and Kawasaki haplotypes were aligned, visually inspected, and compared to previously identified motifs.

### Analyses of structural variants identified in CH17 IGL and IGK BAC clones

Using the program miropeats ^56^ CH17 contigs for IGK and IGL loci were compared individually to the sequences from the Kawasaki haplotypes (IGL, accession NG_000002.1; IGK-proximal/distal, accession NG_000834.1/NG_000833.1). The outputs for each comparison were visually inspected for potential regions of structural variation. Putative breakpoints for the single variant identified were determined by creating a multi-sequence alignment using sequences from the IGL Kawasaki haplotype and novel BAC clone that spanned the regions of the predicted variant-associated breakpoints. Multi-sequence alignments were generated and visualized in SeqMan Pro. Gene prediction was carried out using Genscan (genes.mit.edu/GENSCAN.html; ^57^), following by BLAST using the non-redundant gene database.

### Comparisons of IGK proximal and distal sequence similarity and recombination analysis

For comparisons of proximal and distal V regions of the IGK locus from both the CH17 and Kawasaki haplotypes, only paralogous sequence shared between the proximal and distal regions were considered (*i.e.*, sequence spanning the genes *IGKV1-6*, *IGKV1-5*, *IGKV5-2*, and *IGKV4-1* was excluded, as these genes do have paralogous duplicates in the distal IGKV region). Base pair differences were collated based on pair-wise global alignments made between the Kawasaki proximal sequence and all other proximal and distal sequences from the Kawasaki and CH17 haplotypes. Global alignments and variant calls were carried out using “run-mummer3” and “combineMUMs” commands in MUMmer3.0 ^58^. A sequence similarity plot was then generated for each pair-wise comparison using 10 kbp windows with a sliding size of 500 bp, as reported previously ^17^. Sequences, ∼22.5 kbp in length, from the region suspected to harbor a potential recombination event between proximal and distal regions, were extracted from each haplotype (proximal and distal) and aligned using ClustalW ^59^ within ebioX (http://www.ebioinformatics.org/ebiox/). Recombination/gene conversion analysis based on this alignment was conducted using the Difference of Sums of Squares (DSS) method within TOPALi v2 ^60,61^. This method is based on comparing the branching patterns of two trees constructed using the first and second halves of sequence alignments within a given window of a larger alignment being analyzed; the fit between these two trees and the calculation of DSS is measured using the sum of squares. Windows in which trees differ significantly between the two halves are scored with high DSS values, and are thus candidate sites for recombination. The parameters used here for this analysis were as follows: a window size of 2.5 Kbp with a step size of 100 bp; the Jukes-Cantor substitution model for calculating distance matrices; 500 bootstrap iterations to test for significance; and the analysis was conducted in both forward and reverse directions along the alignment.

### Analysis of IG loci genomic features, locus-wide alignments, and SNP discovery

Locus-wide SNPs were first called in the IGLV, IGKV, and IGHV regions of the CH17 haplotypes by conducting global alignments of the CH17 and Kawasaki or Matsuda haplotypes. For IGHV, sequence 10 kbp downstream of *IGHV6-1* (the most proximal IGHV gene) to 49 kbp upstream of *IGHV3-74* (the most distal IGHV gene) from both CH17 and Matsuda haplotypes ^7,31^ were compared; structural variants identified between the two haplotypes were removed, leaving 834,802 bp of aligned sequence. For IGKV and IGLV, 10 kbp downstream of the most proximal V gene and 10 Kbp upstream of the most distal V gene were compared totalling 858,805 and 858,244 bp of aligned sequence, respectively (the large insertion variant identified in CH17 within IGLV was not included). The lengths of aligned sequence are based on bp coordinates in the Matsuda (IGH) and Kawasaki (IGL and IGK) haplotypes. CH17 and reference haplotypes were aligned on a per locus basis and SNPs were determined from the resulting alignments using the same commands from MUMmer3.0 referenced above. Single nucleotide variants called from CHM1 cell line DNA using whole-genome paired-end Illumina short-read sequencing (Steinberg *et al*.*, manuscript in preparation*; NCBI BioProject ID: 176729) were used to assess the accuracy of variants called from the CH17 assemblies. To do this, the NCBI remap tool (http://www.ncbi.nlm.nih.gov/genome/tools/remap) was used to convert the GRCh37 coordinates to the CHM1_1.1 assembly coordinates (Genbank Accession: GCA_000306695.2), which were then compared with variant calls generated from the alignment of CHM1 Illumina data to the CHM1_1.1 assemblies. Variants in both callsets were flagged as unsupported by the Illumina data, and deemed errors in the CHM1_1.1 assembly.

The coordinates of V gene exon boundaries based on the *Vega* gene annotation track, repeat content (RepeatMasker 3.2.7; www.repeatmasker.org), SDs ^62,63^, and centromere/telomere coordinates were downloaded from UCSC (www.genome.ucsc.edu). Percent sequence identity values for SDs were also downloaded from UCSC and used for partitioning the SD datasets. Locus coverage and SNP density values were calculated using BEDTools version 2.1. We assessed the statistical significance for the observed enrichments of CH17 SNP densities within SDs in the IGHV and IGKV regions using the Genomic Hyperbrowser (https://hyperbrowser.uio.no). We created “case-control” tracks including coordinates overlapped (SD) and not overlapped (non-SD) by SDs in the IGHV/IGKV regions, and then carried out tests for enrichments of CH17 SNPs in these regions by permuting the SD and non-SD status for each IGHV and IGKV set of coordinates (Monte Carlo Simulations = 10,000). The observed enrichments were then compared to the simulated datasets to calculate *P* values for IGHV and IGKV region analyses. For genome-wide centromere/telomere analysis, we used CHM1 variants, and SNP densities in telomeric/centromeric regions were estimated twice independently using telomere/centromere coordinates extended by either 1Mbp or 3Mbp.

## Acknowledgements

We are grateful to Marie-Paule Lefranc and to the IMGT Nomenclature Committee for their help in defining IG genes and alleles. C.T.W. was supported in part by a President’s Research Stipend and graduate fellowship awarded by Simon Fraser University. K.M.S. was supported by a Ruth L. Kirschstein National Research Service Award (NRSA) training grant to the University of Washington (T32HG00035) and an individual NRSA Fellowship (F32GM097807). This work was supported by the US National Institutes of Health (grants 2R01HG002385 and 5P01HG004120 to E.E.E.) and a National Science and Engineering Research Council of Canada grant to F.B. E.E.E. is an Investigator of the Howard Hughes Medical Institute.

## Conflict of Interest Statement

E.E.E. is on the scientific advisory board (SAB) for DNAnexus and was an SAB member of Pacific Biosciences, Inc. (2009-2013) and SynapDx Corp. (2011-2013).

**GenBank Accessions:**

CH17-118P18 AC244205

CH17-436P12 AC243970

CH17-405H5 AC245015

CH17-140P2 AC244255

CH17-158B1 AC233264

CH17-98L16 AC245506

CH17-216K2 AC243981

CH17-68J4 AC247037

CH17-335O14AC245452

CH17-424L4 AC245517

CH17-264L24 AC245060

CH17-238D3 AC245291

CH17-242N13AC246793

CH17-420I24 AC244250

CH17-475O7 AC244157

CH17-95F2 AC245028

CH17-355P2 AC245054

CHM1_1.1 Assembly GCA_000306695.2

IMGT Accessions: submitted to IMGT, waiting for approval

## References

1 Murphy, K., Travers, P. & Walport, M. Janeway’s immunobiology. 7 edn, (Garland Science, 2007).

2 Lefranc, M. P. & Lefranc, G. The immunoglobulin FactsBook. (Academic Press, 2001).

3 Tonegawa, S. Somatic generation of antibody diversity. Nature 302, 575–581 (1983).

4 Boyd, S. D. et al. Individual variation in the germline Ig gene repertoire inferred from variable region gene rearrangements. J Immunol 184, 6986–6992, doi:10.4049/jimmunol.1000445 (2010).

5 Glanville, J. et al. Naive antibody gene-segment frequencies are heritable and unaltered by chronic lymphocyte ablation. Proc Natl Acad Sci U S A 108, 20066–20071, doi:10.1073/pnas.1107498108 (2011).

6 Kidd, M. J. et al. The inference of phased haplotypes for the immunoglobulin H chain V region gene loci by analysis of VDJ gene rearrangements. J Immunol 188, 1333–1340, doi:10.4049/jimmunol.1102097 (2012).

7 Watson, C. T. et al. Complete haplotype sequence of the human immunoglobulin heavy-chain variable, diversity, and joining genes and characterization of allelic and copy-number variation. Am J Hum Genet 92, 530–546 (2013).

8 Watson, C. T. & Breden, F. The immunoglobulin heavy chain locus: genetic variation, missing data, and implications for human disease. Genes and immunity 13, 363–373, doi:10.1038/gene.2012.12 (2012).

9 Ruiz, M., Pallares, N., Contet, V., Barbi, V. & Lefranc, M. P. The human immunoglobulin heavy diversity (IGHD) and joining (IGHJ) segments. Exp Clin Immunogenet 16, 173–184, doi:19109 (1999).

10 Feeney, A. J., Atkinson, M. J., Cowan, M. J., Escuro, G. & Lugo, G. A defective Vkappa A2 allele in Navajos which may play a role in increased susceptibility to haemophilus influenzae type b disease. J Clin Invest 97, 2277–2282 (1996).

11 Sui, J. et al. Structural and functional bases for broad-spectrum neutralization of avian and human influenza A viruses. Nature structural & molecular biology 16, 265–273, doi:10.1038/nsmb.1566 (2009).

12 Tsai, F.-J. et al. Identification of novel susceptibility Loci for kawasaki disease in a Han chinese population by a genome-wide association study. PLoS One 6 (2011).

13 Kawasaki, K. et al. One-megabase sequence analysis of the human immunoglobulin lambda gene locus. Genome Res 7, 250–261 (1997).

14 Ghanem, N. et al. Polymorphism of immunoglobulin lambda constant region genes in populations from France, Lebanon and Tunisia. Exp Clin Immunogenet 5, 186–195 (1988).

15 Lefranc, M. P., Pallares, N. & Frippiat, J. P. Allelic polymorphisms and RFLP in the human immunoglobulin lambda light chain locus. Hum Genet 104, 361–369 (1999).

16 Taub, R. A. et al. Variable amplification of immunoglobulin lambda light-chain genes in human populations. Nature 304, 172–174 (1983).

17 Kawasaki, K. et al. Evolutionary dynamics of the human immunoglobulin kappa locus and the germline repertoire of the Vkappa genes. Eur J Immunol 31, 1017–1028 (2001).

18 Osoegawa, K. et al. A bacterial artificial chromosome library for sequencing the complete human genome. Genome Res 11, 483–496 (2001).

19 Juul, L., Hougs, L. & Barington, T. A new apparently functional IGVK gene (VkLa) present in some individuals only. Immunogenetics 48, 40–46 (1998).

20 Kay, P. H., Moriuchi, J., Ma, P. J. & Saueracker, E. An unusual allelic form of the immunoglobulin lambda constant region genes in the Japanese. Immunogenetics 35, 341–343 (1992).

21 Moraes Junta, C. & Passos, G. A. Genomic EcoRI polymorphism and cosmid sequencing reveal an insertion/deletion and a new IGLV5 allele in the human immunoglobulin lambda variable locus (22q11.2/IGLV). Immunogenetics 55, 10–15, doi:10.1007/s00251-003-0549-x (2003).

22 Frippiat, J. P., Dard, P., Marsh, S., Winter, G. & Lefranc, M. P. Immunoglobulin lambda light chain orphons on human chromosome 8q11.2. Eur J Immunol 27, 1260–1265 (1997).

23 Romo-Gonzalez, T. & Vargas-Madrazo, E. Substitution patterns in alleles of immunoglobulin V genes in humans and mice. Mol Immunol 43, 731–744, doi:10.1016/j.molimm.2005.03.018 (2006).

24 Pallares, N., Lefebvre, S., Contet, V., Matsuda, F. & Lefranc, M. P. The human immunoglobulin heavy variable genes. Exp Clin Immunogenet 16, 36–60 (1999).

25 Genomes Project, C. et al. An integrated map of genetic variation from 1,092 human genomes. Nature 491, 56–65, doi:10.1038/nature11632 (2012).

26 Brensing-Kuppers, J., Zocher, I., Thiebe, R. & Zachau, H. G. The human immunoglobulin kappa locus on yeast artificial chromosomes (YACs). Gene 191, 173–181 (1997).

27 Barbie, V. & Lefranc, M. P. The human immunoglobulin kappa variable (IGKV) genes and joining (IGKJ) segments. Exp Clin Immunogenet 15, 171–183 (1998).

28 Jaenichen, H. R., Pech, M., Lindenmaier, W., Wildgruber, N. & Zachau, H. G. Composite human VK genes and a model of their evolution. Nucleic Acids Res 12, 5249–5263 (1984).

29 Bailey, J. A. & Eichler, E. E. Primate segmental duplications: crucibles of evolution, diversity and disease. Nat Rev Genet 7, 552–564 (2006).

30 Eichler, E. E. Recent duplication, domain accretion and the dynamic mutation of the human genome. Trends Genet 17, 661–669 (2001).

31 Matsuda, F. et al. The complete nucleotide sequence of the human immunoglobulin heavy chain variable region locus. The Journal of experimental medicine 188, 2151–2162 (1998).

32 Arndt, P. F., Hwa, T. & Petrov, D. A. Substantial regional variation in substitution rates in the human genome: importance of GC content, gene density, and telomere-specific effects. J Mol Evol 60, 748–763 (2005).

33 Collins, A. M. et al. The reported germline repertoire of human immunoglobulin kappa chain genes is relatively complete and accurate. Immunogenetics 60, 669–676 (2008).

34 Jackson, K. J. L. et al. Divergent human populations show extensive shared IGK rearrangements in peripheral blood B cells. Immunogenetics 64, 3–14 (2012).

35 Wang, Y. et al. Genomic screening by 454 pyrosequencing identifies a new human IGHV gene and sixteen other new IGHV allelic variants. Immunogenetics 63, 259–265 (2011).

36 Eichler, E. E. Masquerading repeats: paralogous pitfalls of the human genome. Genome Res 8, 758–762 (1998).

37 Estivill, X. et al. Chromosomal regions containing high-density and ambiguously mapped putative single nucleotide polymorphisms (SNPs) correlate with segmental duplications in the human genome. Hum Mol Genet 11, 1987–1995 (2002).

38 Nadel, B. et al. Decreased frequency of rearrangement due to the synergistic effect of nucleotide changes in the heptamer and nonamer of the recombination signal sequence of the V kappa gene A2b, which is associated with increased susceptibility of Navajos to Haemophilus influenzae type b disease. J Immunol 161, 6068–6073 (1998).

39 Romo-Gonzalez, T. & Vargas-Madrazo, E. Structural analysis of substitution patterns in alleles of human immunoglobulin VH genes. Mol Immunol 42, 1085–1097, doi:10.1016/j.molimm.2004.11.004 (2005).

40 Pargent, W., Schable, K. F. & Zachau, H. G. Polymorphisms and haplotypes in the human immunoglobulin kappa locus. Eur J Immunol 21, 1829–1835, doi:10.1002/eji.1830210808 (1991).

41 Schaible, G., Rappold, G. A., Pargent, W. & Zachau, H. G. The immunoglobulin kappa locus: polymorphism and haplotypes of Caucasoid and non-Caucasoid individuals. Hum Genet 91, 261–267 (1993).

42 Sharp, A. J. et al. Segmental duplications and copy-number variation in the human genome. Am J Hum Genet 77, 78–88, doi:10.1086/431652 (2005).

43 Hurles, M. E. Gene conversion homogenizes the CMT1A paralogous repeats. BMC Genomics 2, 11–11 (2001).

44 Hallast, P., Nagirnaja, L., Margus, T. & Laan, M. Segmental duplications and gene conversion: Human luteinizing hormone/chorionic gonadotropin beta gene cluster. Genome Res 15, 1535–1546 (2005).

45 Verrelli, B. C. & Tishkoff, S. A. Signatures of selection and gene conversion associated with human color vision variation. Am J Hum Genet 75, 363–375 (2004).

46 Xu, J. L. & Davis, M. M. Diversity in the CDR3 region of V(H) is sufficient for most antibody specificities. Immunity 13, 37–45 (2000).

47 Avnir, Y. et al. Molecular signatures of hemagglutinin stem-directed heterosubtypic human neutralizing antibodies against influenza A viruses. PLoS Pathog 10 (2014).

48 Tang, X.-C. et al. Identification of human neutralizing antibodies against MERS-CoV and their role in virus adaptive evolution. Proc Natl Acad Sci U S A 111, 2018–2026 (2014).

49 Frippiat, J. P. et al. Organization of the human immunoglobulin lambda light-chain locus on chromosome 22q11.2. Hum Mol Genet 4, 983–991 (1995).

50 Zachau, H. G. The immunoglobulin kappa locus-or-what has been learned from looking closely at one-tenth of a percent of the human genome. Gene 135, 167–173 (1993).

51 Zachau, H. G. The immunoglobulin kappa genes. Immunologist 135, 167–173 (1996).

52 Altschul, S. F., Gish, W., Miller, W., Myers, E. W. & Lipman, D. J. Basic local alignment search tool. J Mol Biol 215, 403–410 (1990).

53 Brochet, X., Lefranc, M. P. & Giudicelli, V. IMGT/V-QUEST: the highly customized and integrated system for IG and TR standardized V-J and V-D-J sequence analysis. Nucleic Acids Res 36, W503–508, doi:10.1093/nar/gkn316 (2008).

54 Giudicelli, V., Brochet, X. & Lefranc, M.-P. IMGT/V-QUEST: IMGT standardized analysis of the immunoglobulin (IG) and T cell receptor (TR) nucleotide sequences. Cold Spring Harb Protoc 2011, 695–715 (2011).

55 Quinlan, A. R. & Hall, I. M. BEDTools: a flexible suite of utilities for comparing genomic features. Bioinformatics 26, 841–842, doi:10.1093/bioinformatics/btq033 (2010).

56 Parsons, J. D. Miropeats: graphical DNA sequence comparisons. Comput Appl Biosci 11, 615–619 (1995).

57 Burge, C. & Karlin, S. Prediction of complete gene structures in human genomic DNA. J Mol Biol 268, 78–94 (1997).

58 Kurtz, S. et al. Versatile and open software for comparing large genomes. Genome Biol 5 (2004).

59 Thompson, J. D., Gibson, T. J. & Higgins, D. G. Multiple sequence alignment using ClustalW and ClustalX. Curr Protoc Bioinformatics Chapter 2 (2002).

60 McGuire, G. & Wright, F. TOPAL 2.0: improved detection of mosaic sequences within multiple alignments. Bioinformatics 16, 130–134 (2000).

61 Milne, I. et al. TOPALi: software for automatic identification of recombinant sequences within DNA multiple alignments. Bioinformatics 20, 1806–1807, doi:10.1093/bioinformatics/bth155 (2004).

62 Bailey, J. A., Yavor, A. M., Massa, H. F., Trask, B. J. & Eichler, E. E. Segmental duplications: organization and impact within the current human genome project assembly. Genome Res 11, 1005–1017, doi:10.1101/gr.187101 (2001).

63 Bailey, J. A. et al. Recent segmental duplications in the human genome. Science 297, 1003–1007, doi:10.1126/science.1072047 (2002).

